# Application of the Nicking Loop™ targeted library preparation method to DNBSEQ™ sequencing

**DOI:** 10.64898/2026.03.10.710732

**Authors:** Simona Adamusová, Anttoni Korkiakoski, Tatu Hirvonen, Hui Ren, Nea Laine, Anna Musku, Tuula Rantasalo, Jorma Kim, Juuso Blomster, Jukka Laine, Chongjun Xu, Manu Tamminen, Juha-Pekka Pursiheimo

## Abstract

Nicking Loop™ is a PCR-free targeted library preparation method that combines conversion of linear target DNA into circular single-stranded DNA (CssDNA) library with early sample indexing in a single step. The resulting CssDNA libraries can be either directly sequenced or optionally amplified, offering maximum flexibility across sequencing applications.

This study demonstrates the compatibility of Nicking Loop™ circular libraries with a MGI’s DNBSEQ™ platform. Compatibility was evaluated against established linear Nicking Loop™ libraries sequenced on Illumina MiSeq platform. Using synthetic reference samples with defined variant allele frequencies, Nicking Loop™ method demonstrated matching performance across both library formats and sequencing platforms. Key quality metrics, including unique molecular identifier (UMI) distributions, error profiles and VAF detection, were all highly consistent. Both library types generated over 97% singleton UMIs, indicating uniform template sampling, and VAF measurements were strongly concordant across platforms (Spearman’s ρ = 0.939). Collectively, these findings demonstrate that Nicking Loop™ method is directly applicable to circular NGS platforms, such as DNBSEQ™, strongly supporting its use as a platform-agnostic library preparation strategy for targeted sequencing applications.

## INTRODUCTION

The field of next-generation sequencing (NGS) expanded rapidly over the past decade. In addition to established short-read sequencing platforms such as Illumina, technologies developed by MGI, Element Biosciences, and Pacific Biosciences introduced alternative sequencing chemistries, including the use of circular templates and long-read sequencing strategies^1–3^. However, many library preparation workflows are optimized for linear, Illumina-like libraries and often require additional conversion steps to generate libraries compatible with circular approaches^4^. In addition, library preparation protocols for newer NGS platforms are commonly optimized for whole-genome sequencing applications primarily used in research ^5–7^. In contrast, targeted sequencing approaches remain widely used in clinical and diagnostic settings due to cost-efficiency, high-depth coverage, and assay design considerations^8,9^.

Nicking Loop™ is a targeted library preparation method that converts linear DNA into circular single-stranded DNA (CssDNA) while introducing sample-specific index at the early step of the workflow^10^. Early sample indexing enables direct sequencing of circular libraries following conversion or pooling of samples prior to further library amplification if desired.

Nicking Loop™ leverages linear amplification to limit error propagation, which is a key attribute of circular NGS platforms. These features make Nicking Loop™ well suited for sequencing platforms that utilize circular library formats. This compatibility is particularly relevant for applications where detection of low-frequency variants is critical, including detection of somatic variants from circulating cell-free DNA or tissue, and infectious disease testing.

The Nicking Loop™ has been previously evaluated on multiple sequencing platforms, and this study extends the evaluation to MGI’s DNBSEQ™ platform. Here, the performance of circular Nicking Loop™ libraries sequenced on DNBSEQ™-G99 is compared to linear Nicking Loop™ libraries sequenced on Illumina MiSeq. Platform compatibility is assessed using key performance metrics, including unique molecular identifier distributions, error profiles and variant allele frequency (VAF) profiling capabilities, with early sample indices used for read assignment on DNBSEQ™ platform.

## MATERIAL AND METHODS

### Oligonucleotides

The components of a Nicking Loop construct: the Loop carrying the sample index, left and right probes, and the bridge oligonucleotide and two synthetic DNA pools used as target DNA were produced by IDT (Coralville, IA). Left and right probes were pre-annealed with the bridge oligonucleotide to form a probe construct. The DNA pools contained ten templates. Individual templates were distinguished by three pool-specific complementary nucleotides. The pools were mixed in different ratios to mimic the VAF of 0%, 1%, 5%, 10%, and 20%.

### Preparation of Nicking Loop™ converted CssDNA and RCA amplification

To prepare Nicking Loop™ converted CssDNA from linear target DNA, 2.5 fmol of linear synthetic target pools were combined with 10 fmol of probe constructs (probes 6 & 7 were loaded at 1 fmol instead) and 200 fmol of Loop oligomer in 1× Ampligase buffer (LGC Biosearch Technologies, Hoddesdon, UK). To induce hybridization, the mixture was denatured at 95°C for 5 minutes, cooled stepwise to 55°C (10°C steps with 2.5 minutes hold), and incubated at 55°C for 2 hours. To circularize the hybridization products, the reactions were supplemented with: 0.2 U Phusion High-Fidelity DNA Polymerase (Thermo Fisher Scientific, Waltham, MA), 0.2 U Ampligase (LGC Biosearch Technologies), and 10 mM dNTPs (Thermo Fisher Scientific) in 1 × Ampligase Reaction buffer (LGC Biosearch Technologies) and incubated at 55°C for 40 minutes.

To degrade residual linear DNA, 1 µL of each Thermolabile Exonuclease I and RecJf (NEB, Ipswich, MA) was added to the reaction, and incubated at 37°C for 45 minutes, followed by heat inactivation at 80°C for 10 minutes. Nicking Loop™ converted CssDNA templates were purified using a 2× volume of AMPure XP beads (Beckman Coulter, Brea, CA).

Nicking Loop™ converted CssDNA templates prepared by conversion of linear DNA were amplified by rolling-circle amplification (RCA). In the first step, 10 µL of purified CssDNA templates were pre-annealed with 12.5 pmol of universal amplification primer in 1× EquiPhi29 buffer (Thermo Fisher Scientific). The mixture was denatured at 95°C for 5 minutes and gradually cooled down to room temperature (RT) for 20 minutes. To produce RCA concatemers, the reaction was supplemented with 10 mM dNTPs, 100 mM DTT, and 0.5 U of EquiPhi29 (Thermo Fisher Scientific) followed by incubation at 45°C for 90 minutes, and subsequent heat inactivation at 95°C for 10 minutes.

### Circular library preparation for DNBSEQ™-99

The circular Nicking Loop™ libraries were prepared from RCA-generated concatemers. To promote folding of the concatemers, reactions were supplemented with nuclease-free water and 1× rCutSmart buffer (NEB), incubated at 95°C for 5 minutes, and gradually cooled to RT. Folded concatemers formed a hairpin structure with a dsDNA restriction recognition site for the nicking enzyme. To produce linear monomers, the recognition site was digested with 10 U of Nb.BsrDI (NEB) or 15 U of Nb.BbvCI (NEB) at 37°C for 90 minutes, followed by heat inactivation at 80°C for 10 minutes. The resulting linear monomers were purified using 1.8× volume of AMPure XP beads (Beckman Coulter).

To initiate folding of monomers into a circular Nicking Loop structure with a nick, 10 µL of purified linear monomers was mixed with 7 µL of nuclease-free water, denatured at 95°C for 5 minutes, and gradually cooled to RT. The nick was subsequently ligated by addition of 1 µL of T4 DNA ligase and 2 µL of 10× T4 DNA ligase buffer (NEB), followed by incubation at 32°C for 30 minutes. Any residual linear ssDNA was degraded by 1 µL each of Thermolabile Exonuclease I and RecJf (NEB). The reactions were incubated at 37°C for 45 minutes, then heat-inactivated at 95°C for 10 minutes. The final circular library was purified using a 2× volume of AMPure XP beads (Beckman Coulter). The libraries were quantified using the Qubit™ dsDNA HS Assay Kit and Qubit™ 4 Fluorometer (Thermo Fisher Scientific). A schematic of the circular Nicking Loop™ library is provided in Supplementary Figure. 1.

**Figure 1.**
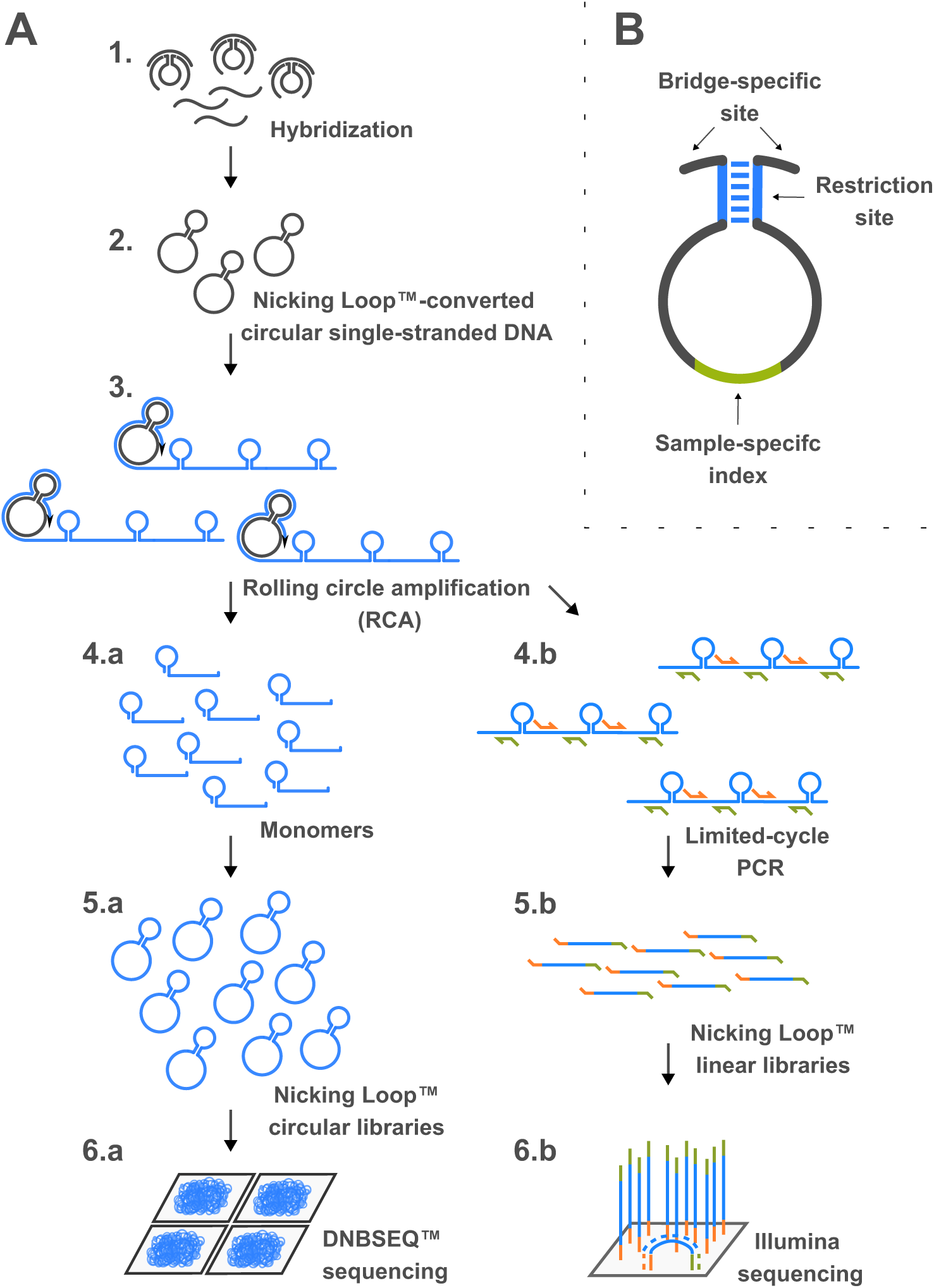
Nicking Loop™ targeted library preparation method. **(A)** The conversion of linear DNA to circular DNA and platform-specific processing steps for circular DNBSEQ™ and linear Illumina platform. The Nicking Loop™ construct comprises a bridge oligonucleotide, the Loop oligonucleotide, and left and right target-specific probes that bind to target DNA (1). Hybridized construct is circularized, forming Nicking Loop™-converted circular single-stranded DNA (CssDNA; 2) that is amplified by rolling circle amplification (RCA). The RCA-amplified concatemers are used to prepare both circular and linear libraries (3). For circular libraries, the concatemers are cleaved into monomers (4a). The monomers fold and circularize, generating circular libraries (5a) that are sequenced with DNBSEQ™ platform (6a). For linear libraries, RCA amplified concatemers are subjected to limited-cycle PCR (4b) to generate linear libraries (5b) that are sequenced with Illumina platform (6b). **(B)** The Loop oligonucleotide consists of bridge-specific site required for incorporation into hybridized construct, complementary restriction site for nicking endonuclease, and sample-specific index used for early sample indexing.

### DNBSEQ™-G99 Sequencing

Nicking Loop™ libraries were amplified into DNA nanoballs (DNBs) and sequenced on the DNBSEQ™-G99 platform following the manufacturer’s system guide (Complete Genomics, San Jose, CA). Briefly, for DNB preparation, 250 fmol of pooled library was combined with 10 μL of Make DNB Buffer containing primer for RCA amplification. Then appropriate amount of TE buffer was added to make the final volume of 20 μL for primer hybridization. Hybridization was performed with sequential incubations at 95 °C, 65 °C, and 40 °C for 1 minute each. Subsequently, 20 μL of Make DNB Enzyme Mix I and 2 μL of Make DNB Enzyme Mix II (LC) were added, and DNB amplification was carried out at 30 °C for 30 minutes. The reaction was terminated by adding 10 μL of Stop DNB Reaction Buffer. DNB concentration was quantified using the Qubit ssDNA Assay Kit (Thermo Fisher Scientific, PN #Q10212) by using 2 μL of above DNB solution. Then, 21 μL of DNB solution was mixed with 7 μL of DNB Load Buffer II and 1 μL of Make DNB Enzyme Mix II (LC), then loaded onto the flow cell according to manufacturer’s system guide. Sequencing was performed using the DNBSEQ™-G99RS High-Throughput Sequencing Set (PN #940-001717-00, Model APP-D FCL PE300, Version 1.0). Run parameters were configured for paired-end sequencing with Read 1 of 66 bp, Read 2 of 121 bp, and a 45 bp barcode read.

Custom-designed Make DNB primers, sequencing primers, and MDA primers based on the Nicking Loop™ library structure were used for DNB amplification and library sequencing (Supplementary Fig. 1).

### Library preparation and sequencing with Illumina MiSeq

The linear Nicking Loop™ libraries were produced from RCA-generated concatemers (see section *Preparation of Nicking Loop™ converted CssDNA and RCA amplification*). To introduce flowcell binding sequences, 2 µL of RCA concatemers was subjected to minimal-cycle PCR using 10 µM dual-indexed primers with Phusion HS II DNA Polymerase (Thermo Fisher Scientific), following the manufacturer’s guidelines. PCR was performed with a combined annealing and extension step at 72°C for 14 cycles. All PCR products were gel-purified using the Monarch DNA Gel Extraction kit (NEB). The linear libraries were quantified using the Qubit™ dsDNA HS Assay Kit and the Qubit™ 4 Fluorometer (Thermo Fisher Scientific). Sequencing was performed on the Illumina MiSeq platform, using the MiSeq Reagent Kit v3 for a 2 × 300 bp paired-end run (Illumina). The schematic of the linear Nicking Loop™ library structure is provided in Supplementary Figure 2.

### Data analysis

Reads generated by DNBSEQ™ platform and Illumina MiSeq were merged using VSEARCH ^11^ (v2.15.2_linux_x86_64) with the following parameters: --fastq_minovlen 10 --fastq_maxdiffs 15 --fastq_maxee 1 --fastq_allowmergestagger --fastq_qmaxout 92 --fastq_qmax 55. Since the sequencing depth of the DNBSEQ™ platform was approximately 225-fold higher (109,080,339 merged reads) than that of the Illumina MiSeq (485,342 merged reads), the merged DNBSEQ™ read pool was subsampled down to 485,342 reads.

The merged reads were discarded if the gap sequence (target region between probes) did not match the target-specific probe sites, had an incorrect gap length, or had discrepancies in three pool-specific nucleotides (2.7% of Illumina MiSeq merged reads and 4.5% of DNBSEQ™-G99 merged reads). The error rate for each platform was determined by comparing the gap sequence to the expected reference sequence.

All data were processed using proprietary pipelines. Python version 3.11.14 (https://www.python.org/downloads/release/python-31114/) and matplotlib library version 3.10.8 (https://matplotlib.org/3.10.8) were used to draw the figures. SciPy v1.17.0 (https://pypi.org/project/scipy/1.17.0/) library was used for Spearman correlations.

## RESULTS

### Nicking Loop™ libraries for circular and linear NGS platforms

To assess compatibility with circular NGS platforms, circular Nicking Loop™ libraries were sequenced on the DNBSEQ™-G99. Nicking Loop™ was initially established using linear libraries on Illumina MiSeq and therefore served as a benchmark for performance of the circular libraries.

As the circular Nicking Loop™ method has been described in detail in a previous study^10^, only the key differences relevant to platform compatibility are summarized here. In the first step, the bridge oligonucleotide, probes, and the Loop are hybridized to linear target-specific DNA. This hybridized intermediate is circularized, forming Nicking Loop™-converted CssDNA that is further amplified by RCA to generate a long concatemer (Fig. 1A, steps 1-3). To prepare a circular library, the concatemers are folded into a hairpin structure with a nicking restriction site in the duplex stem, which is cleaved to produce monomers. The monomers fold to create a circular molecule with a hairpin structure, forming a nick in the stem. This nick is ligated, generating a circular library that can be used with different circular sequencing platforms. For sequencing with the DNBSEQ™ platform, the libraries additionally undergo DNA nanoball generation (Fig 1A, steps 4a–7a).

To prepare libraries for the Illumina platform, the RCA-generated concatemers were subjected to limited-cycle PCR to introduce flowcell binding sequences and produce short linear libraries compatible with bridge amplification (Fig 1A, steps 4b–7b).

### Early sample indexing with Nicking Loop™

The Loop oligonucleotide is an integral component of the Nicking Loop™ library preparation and contains a sample-specific index (Fig. 1B). Upon incorporation of the Loop into the hybridization construct with target DNA, the samples are tagged at an early stage of library preparation. In this study, the early sample-specific index was used to demultiplex sequencing reads generated by the DNBSEQ™-G99 (Supplementary Fig. 3), as the entire structure of the Nicking Loop™ circular library was sequenced (Fig. 1A, step 5a; Supplementary Fig. 1). In contrast, for linear libraries sequenced on the Illumina MiSeq, sample-specific indices were introduced in the final step of library preparation workflow, as the Loop structure was not sequenced by the Illumina MiSeq platform (Fig. 1A, steps 5b; Supplementary Fig. 2).

### UMI distribution across different Nicking Loop™ libraries and NGS platforms

The Nicking Loop™ design incorporates dual UMIs located within the left and right probe arms flanking the target-specific region (Supplementary Fig 1). UMI distributions were assessed across different Nicking Loop™ libraries and NGS platforms. To ensure direct comparability, merged reads of the DNBSEQ™-G99 were subsampled to match the sequencing depth of Illumina MiSeq (approximately 225-fold difference). Despite varying sequencing depths, the general profiles of read and UMI coverage per probe were comparable (Supplementary Fig. 3).

The total UMI count was 429,216 and 459,100 for the DNBSEQ™-G99 (subsampled) and Illumina MiSeq, respectively. Singleton UMIs accounted for 97.36% and 97.03% of total UMIs for the DNBSEQ™™-G99 and the Illumina MiSeq. The UMI distributions for each probe were comparable across platforms (Fig. 2). In some cases, circular Nicking Loop™ libraries sequenced on the DNBSEQ™-G99 platform yielded higher reads per UMI than linear libraries sequenced on Illumina MiSeq, however, these differences were negligible (0.099% of cases).

**Figure 2.**
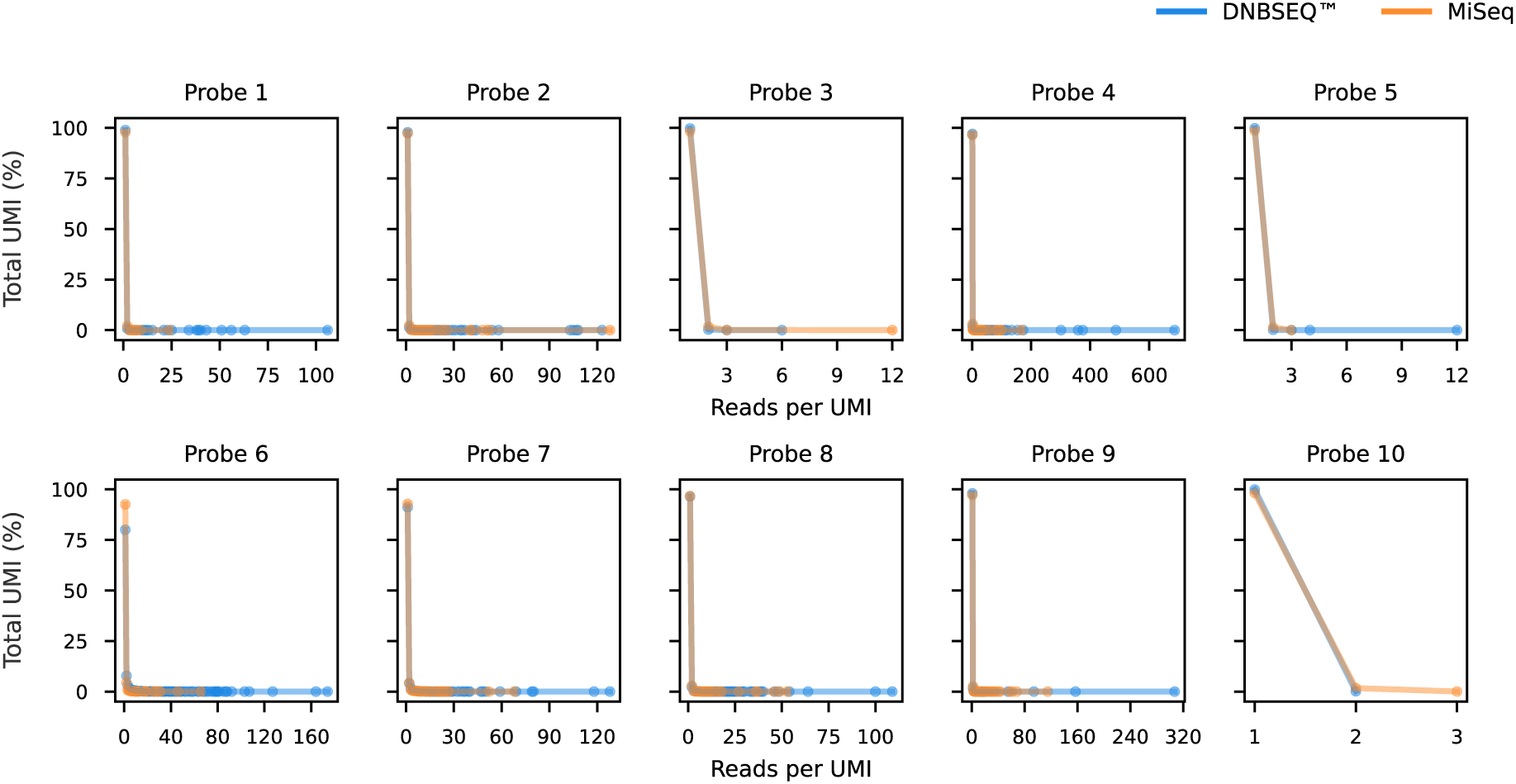
Distribution of reads per unique molecular identifier (UMI) across Nicking Loop™ circular libraries sequenced on the DNBSEQ™-G99 (blue) and linear libraries sequenced on the Illumina MiSeq (orange) for each probe in the synthetic panel.

Beyond the matched-depth comparison of DNBSEQ™-G99, the UMI distribution was assessed at native sequencing depth (109,080,339 merged reads). The distribution remained dominated by UMIs with low read coverage (58.87% singleton UMIs), indicating high molecular diversity (Supplementary Fig. 4).

### Mismatches introduced across different Nicking Loop™ libraries and NGS platforms

To determine the impact of platform-specific Nicking Loop™ steps and sequencing chemistries on error rates, mismatches between the sequenced gap region and the expected reference sequence were evaluated. Error rates were compared using depth-normalized merged reads to ensure direct comparability between platforms.

Overall, fewer mismatches were observed for the circular Nicking Loop™ workflow sequenced on the DNBSEQ™-G99 (99.74% correct reads) compared to linear Nicking Loop™ workflow sequenced on the Illumina MiSeq platform (99.29% correct reads), although the observed difference was small. Single base pair mismatches accounted for the majority of all mismatches (0.25% and 0.70% of reads for DNBSEQ™-G99 and Illumina MiSeq, respectively), while reads containing two or more mismatching base pairs were rare. The error rate of DNBSEQ™-G99 at native sequencing depth was matching the depth-normalized results. At native sequencing depth, 99.76% of reads were correct, 0.23% of reads contained a single base-pair mismatch, and the number of reads with ≥ 2 base-pair mismatches was negligible (Supplementary table 1).

**Table 1.**
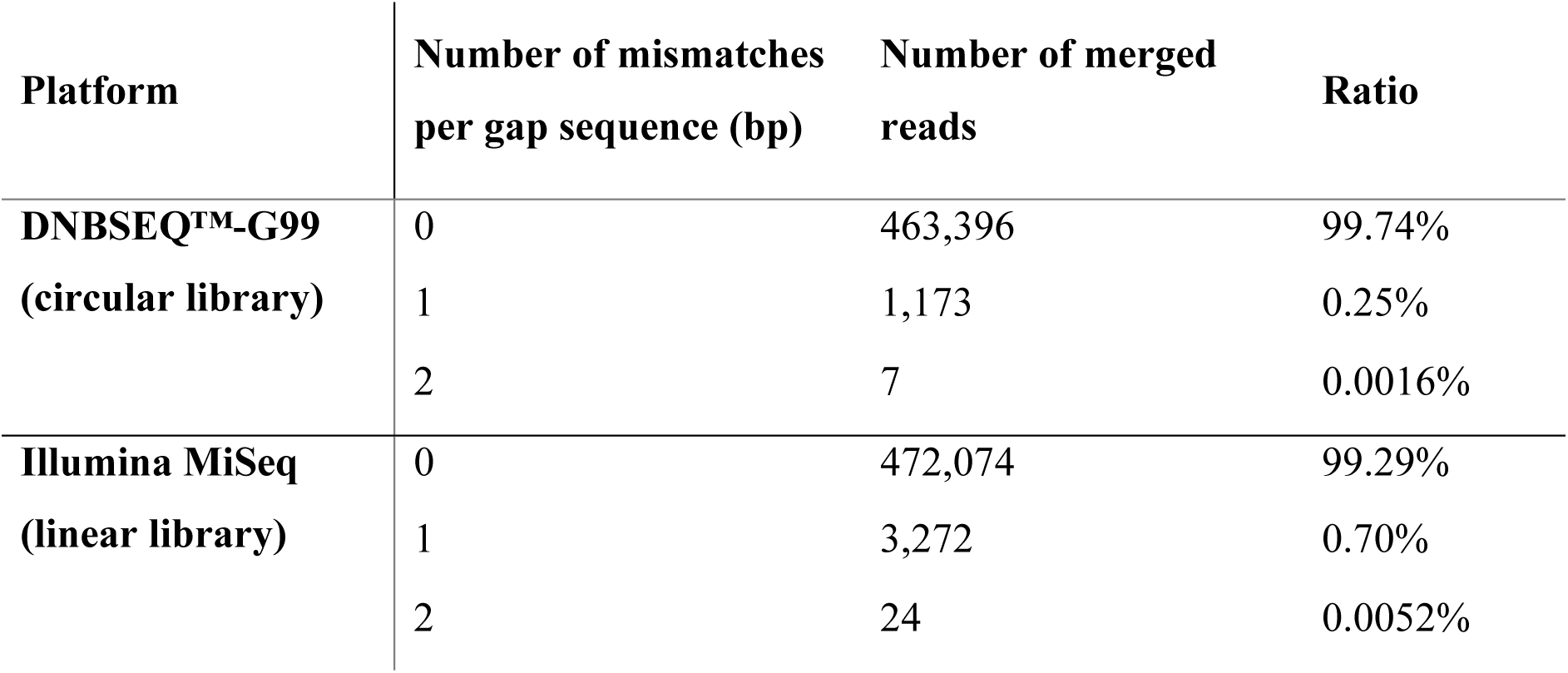
Number of mismatches (bp) observed in gap sequence of merged reads across circular and linear Nicking Loop™ workflows when compared to the expected reference gap sequence. Circular library was sequenced by DNBSEQ™-G99 and linear library was sequenced with Illumina MiSeq. Reads obtained by DNBSEQ™-G99 were subsampled without replacement to match the sequencing depth of Illumina MiSeq.

| Platform | Number of mismatches<br>per gap sequence (bp) | Number of merged<br>reads | Ratio |
| --- | --- | --- | --- |
| **DNBSEQ™-G99**<br>**(circular library)** | 0 | 463,396 | 99.74% |
|  | 1 | 1,173 | 0.25% |
|  | 2 | 7 | 0.0016% |
| **Illumina MiSeq**<br>**(linear library)** | 0 | 472,074 | 99.29% |
|  | 1 | 3,272 | 0.70% |
|  | 2 | 24 | 0.0052% |

### VAF detection across different Nicking Loop™ libraries and NGS platforms

To assess consistency of VAF quantification across library preparation methods and sequencing platforms, synthetic templates mimicking multiple VAFs were processed by Nicking Loop™ and sequenced (Fig. 3). Sequencing results revealed a strong VAF agreement for circular libraries sequenced on DNBSEQ™-G99 and linear libraries sequenced on Illumina MiSeq (Spearman’s ρ = 0.939). The concordance indicates that platform-specific downstream processing steps in the Nicking Loop™ workflow does not affect VAF quantification.

**Figure 3.**
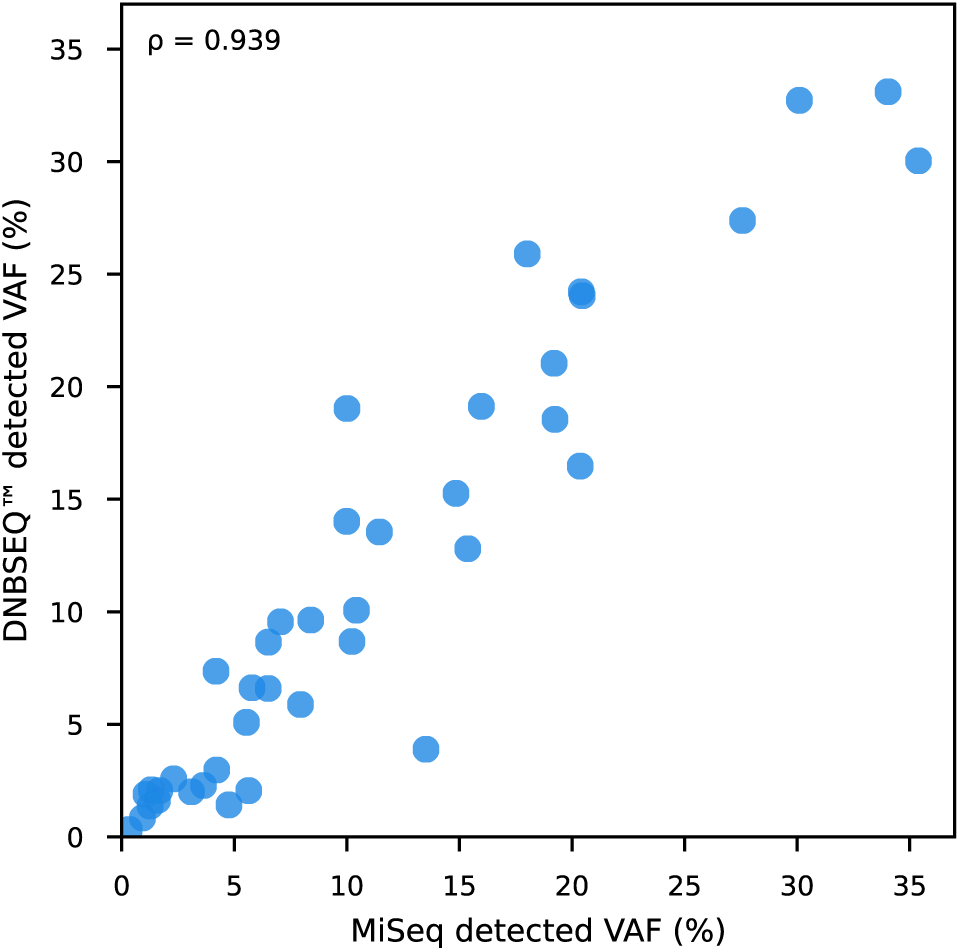
Concordance of variant allele frequencies detected by Nicking Loop™ library preparation workflow for the DNBSEQ™ platform and the Illumina MiSeq. A strong agreement in VAF concordance was observed (Spearman’s ρ = 0.939) despite platform-specific steps of Nicking Loop™ library preparation workflow.

## DISCUSSION

Nicking Loop™ library preparation method provides flexibility for circular NGS platforms by supporting sequencing of circular templates directly converted from linear target DNA or following amplification. While direct circular sequencing offers a simplified protocol relative to traditional library preparation, the amplified Nicking Loop™ libraries offer PCR-free amplification with minimal bias. This study demonstrates that such amplified circular libraries can be effectively sequenced on a circular NGS platforms and demultiplexed using Loop index introduced in the initial step of the workflow, as illustrated by the results from the DNBSEQ™ platform. Furthermore, the agreement observed between circular and linear Nicking Loop™ libraries highlights the robustness and platform-independent read-out of the method, expanding prior demonstrations on PacBio ONSO and Oxford Nanopore platforms^10^. The robustness of the Nicking Loop™ method across library types and NGS platforms was demonstrated by consistent UMI distributions, noise profiles, and reliable VAF detection.

UMI distributions across library types were highly comparable, with both libraries generating more than 97% of singleton UMIs. Negligible deviations were observed, where the circular library exhibited more reads per UMI than linear library, representing less than 0.1% of total UMIs. This effect is likely caused by stochastic amplification of minor number of templates in RCA, and it did not impact VAF detection.

The error profiles observed across the libraries were comparable, indicating that platform-specific steps and sequencing chemistries have a negligible impact on the error rates. Circular libraries sequenced on DNBSEQ™-G99 exhibited fewer errors, however, these differences were within expected variability. Consistent with the observations, the VAFs detected across circular and linear libraries were highly concordant (Spearman’s ρ = 0.939), further supporting the robustness of the method.

Together, these results demonstrate that Nicking Loop™ provides a robust and reproducible platform-independent read-out across multiple NGS platforms. It enables a native targeted sequencing library preparation for circular DNBSEQ™. DNBSEQ™ workflows are primarily optimized for whole-genome sequencing, and commonly used targeted protocols often rely on conversion of PCR-enriched libraries originally designed for linear platforms such as Illumina. In contrast, the Nicking Loop™ library is inherently compatible with circular sequencing architecture, as it directly converts linear target DNA into circular template. The method leverages RCA amplification to limit error propagation, workflow complexity, and material loss associated with library conversion. Importantly, the Nicking Loop™ supports targeted sequencing applications central to clinical use, including high-sensitivity somatic variant detection in oncology and focused gene panels for hereditary disease-testing.

## DATA AVAILABILITY

The data supporting the findings of this study are available upon reasonable request.

## SUPPLEMENTARY DATA

Supplementary Data are available online.

## AUTHOR CONTRIBUTIONS

S.A. – Conceptualization, Methodology, Formal analysis, Writing - Original Draft, Visualization

A.K. – Conceptualization, Methodology, Software, Formal analysis, Data Curation, Writing - Review & Editing, Visualization

T.H. – Conceptualization, Methodology, Writing - Review & Editing, Project administration

H.R. – Methodology, Investigation, Writing - Original Draft, Writing - Review & Editing,

N.L. – Methodology, Software, Formal analysis, Data Curation, Writing - Review & Editing

A.M. – Investigation, Writing - Review & Editing

T.R. – Investigation, Writing - Review & Editing

J.K. – Data Curation, Writing - Review & Editing

J.B. – Writing - Review & Editing

J.L. – Writing - Review & Editing

C.X. – Supervision, Writing - Review & Editing,

M.T. – Conceptualization, Methodology, Supervision, Project administration, Funding acquisition, Writing - Review & Editing

J.P.P. – Conceptualization, Methodology, Investigation, Writing - Original Draft, Supervision, Project administration

## ACKNOWLEDGEMENTS

The authors would like to express their gratitude to Voima Ventures, (Helsinki, Finland), Almaral (Kaarina, Finland), Avohoidon Tutkimussäätiö (Espoo, Finland) and Business Finland (Helsinki, Finland) for support and funding.

## DECLARATION OF GENERATIVE AI AND AI-ASSISTED TECHNOLOGIES IN THE WRITING PROCESS

During the preparation of this work, the authors used ChatGPT 5.2 in order to refine the language and improve readability in certain parts of the manuscript. After using this tool, the authors reviewed and edited the content as needed and take full responsibility for the content of the publication.

## ABBREVIATIONS

CssDNA: Circular single-stranded DNA
DNB: DNA nanoball
dsDNA: Double-stranded DNA
NGS: Next-generation sequencing
ssDNA: Single-stranded DNA
RCA: Rolling-circle amplification
RT: Room temperature
UMI: Unique molecular identifier
VAF: Variant allele frequency

