## Supplementary for "Application of the Nicking Loop™ targeted library preparation method to DNBSEQ™ sequencing"

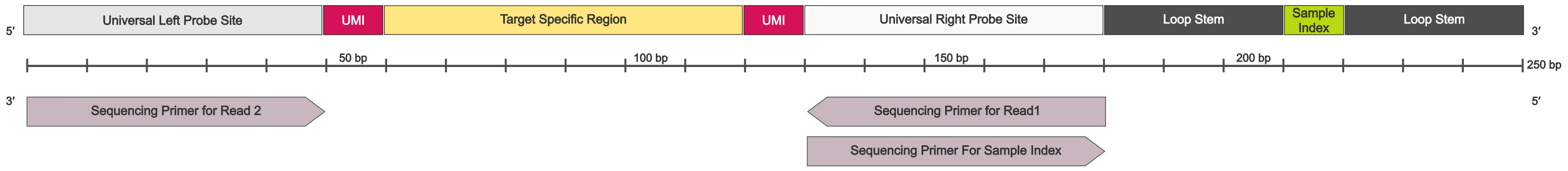


**Supplementary Figure 1.** Circular Nicking Loop™ sequencing library structure for DNBSEQ™ platform. The full length of the library is approximately 250 bp. The library comprises universal left (light grey) and right probe sites (white), unique molecular identifiers (UMIs; pink) flanking the target-specific region (yellow), the Loop stem (dark grey) and the sample index (green). The sample index is introduced in the first step of the workflow. The sequencing primer for read 1, and sequencing primer for the sample index are complementary to the universal right probe site. The sequencing primer for the read 2 is complementary to universal left probe site. The Make DNB and MDA primers (not shown), used for making DNA nanoballs amplification anneal to the universal left probe site.

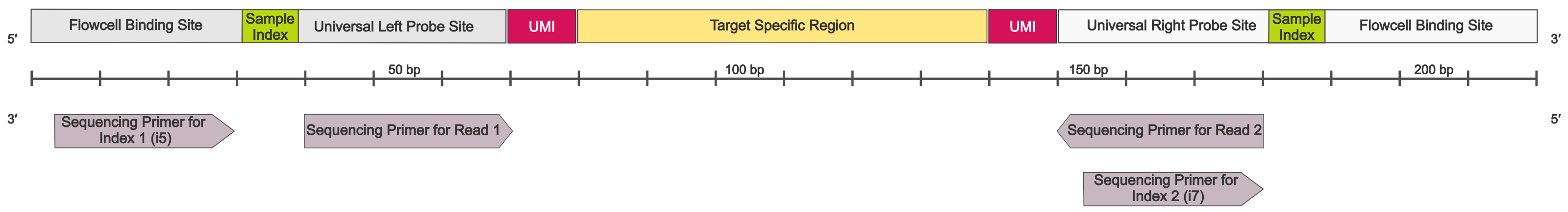


**Supplementary Figure 2.** Linear Nicking Loop™ sequencing library structure for the Illumina platform. The library lenght is approximately 220 bp. It comprises universal left (light grey) and right (white) probe sites with a flowcell binding site, embedded sample index (green), and unique molecular identifiers (UMIs; pink) flanking the target-specific region (yellow). The sample index is introduced during the final indexing PCR step as a part of the indexing primers.

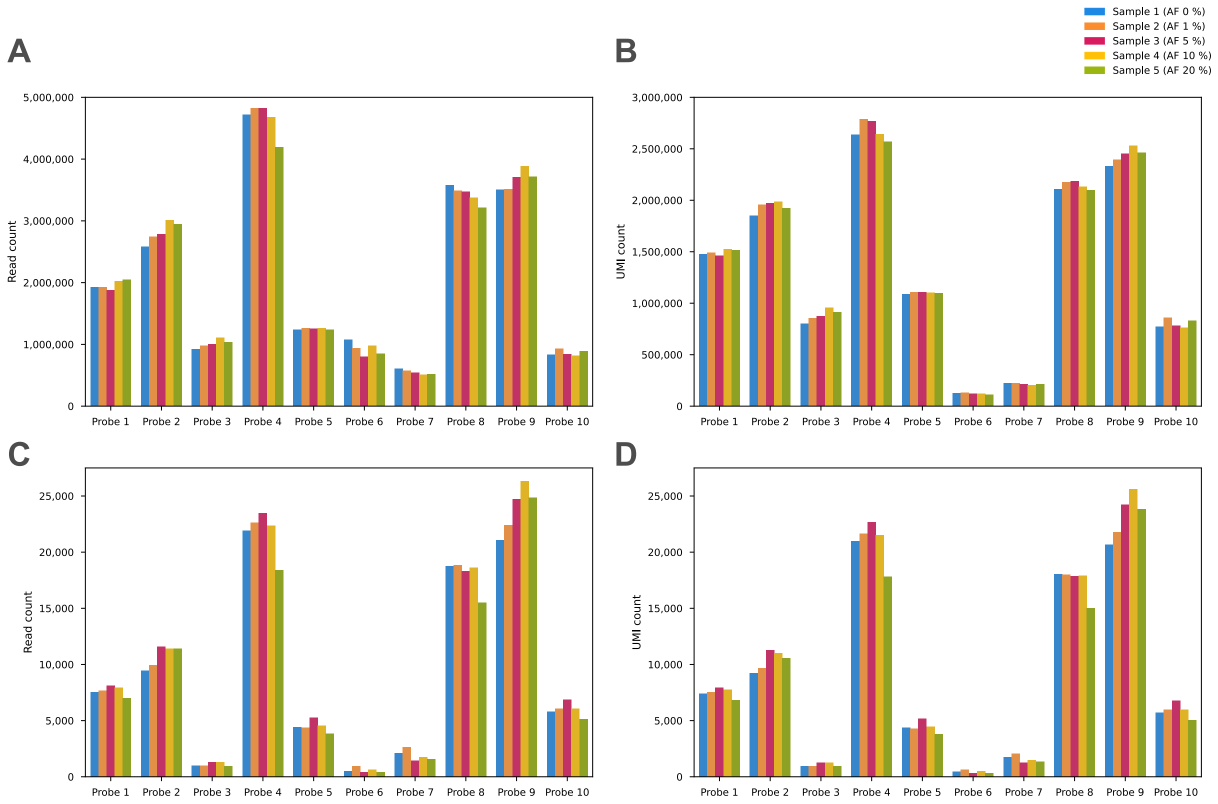


**Supplementary Figure 3.** The sequencing meta-data. The total read count (A) and unique molecular identifier (UMI) count (B) for DNBSEQ™ platform. The total read count (C) and UMI count (D) for Illumina MiSeq. The read and UMI counts are shown for each probe in each sample (indicated by different colour). For DNBSEQ™ platform, the sample reads were demultiplex using the early-sample index from the Loop.

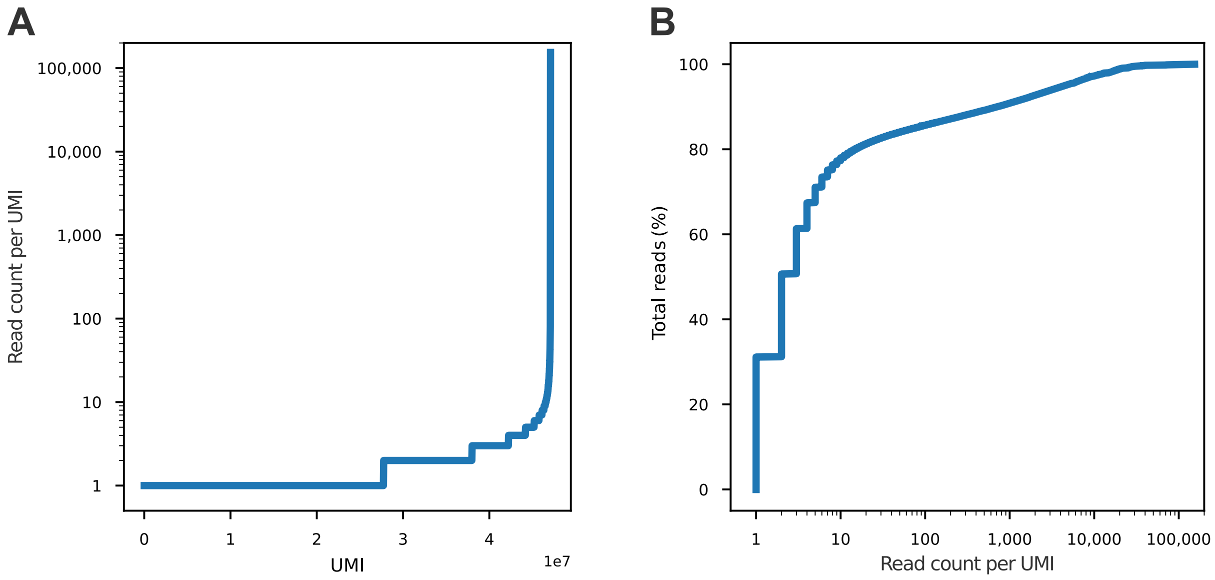


**Supplementary Figure 4**. Distribution of unique molecular identifiers (UMIs) at the native sequencing depth of DNBSEQ-G99. The figure illustrates UMI coverage heterogeneity, and the contribution of UMIs with varying coverage to total sequencing depth. **(A)** Read count per UMI (log scale) plotted against individual UMI (ranked by coverage). **(B)** Cumulative percentage of total reads by read count per UMI (log scale).

**Supplementary Table 1** Number of mismatches (bp) observed in merged reads across circular and linear Nicking Loop™, when compared to the expected reference sequence. Circular library was sequenced by DNBSEQ-G99 and linear library was sequenced with Illumina MiSeq. The DNBSEQ-G99 sequencing resulted in approximately 225-fold more reads than Illumina MiSeq.

| Platform | Number of mismatches per sequence (bp) | Number of merged reads | Ratio |
|---|---|---|---|
| **DNBSEQ-G99 (circular library)** | 0 | 104,161,052 | 99.76% |
| | 1 | 244,558 | 0.23% |
| | 2 | 892 | $8.5 \times 10^{-6}$ % |
| | 3 | 98 | $9.4 \times 10^{-7}$ % |
| | 4 | 9 | $8.6 \times 10^{-8}$ % |
| | 5 | 3 | $2.9 \times 10^{-8}$ % |
| | 6 | 1 | $9.6 \times 10^{-9}$ % |
| **Illumina Miseq (linear library)** | 0 | 472,074 | 99.29% |
| | 1 | 3,272 | 0.70% |
| | 2 | 24 | $5.2 \times 10^{-3}$ % |
